# AhNGE: A Database for *Arachis hypogaea* Nodule Developmental Gene Expression

**DOI:** 10.1101/2021.01.30.428929

**Authors:** Tarannum Shaheen, Kunal Tembhare, Ajeet Singh, Bikash Raul, Asim Kumar Ghosh, Ivone Torres-Jerez, Josh Clevenger, Michael Udvardi, Brian E. Scheffler, Peggy Ozias-Akins, Kaustav Bandyopadhyay, Shailesh Kumar, Senjuti Sinharoy

## Abstract

Symbiotic nitrogen fixation (SNF) inside root-nodules is a primary and sustainable source of soil nitrogen. Understanding nodule development and metabolism in crop legumes may lead to more effective SNF in agriculture. Peanut (*Arachis hypogaea*) is an economically important allotetraploid legume with non-canonical nodule developmental features. Recent genome sequencing of peanut has opened the possibility of making peanut a model for studying atypical nodule development. To help the community of nodule biologists, we have developed a database called AhNGE (*Arachis hypogaea* Nodule Developmental Gene Expression: http://nipgr.ac.in/AhNGE/index.php). AhNGE contains RNAseq data from six data points of nodule development in *A. hypogagea cv. Tifrunner*. This data represents a dynamic view of gene expression during peanut nodule development. Research in model legumes has generated a huge knowledgebase in the last twenty years. To streamline comparative genomics among legumes, we performed ortholog analysis among four legumes (Cicer, Glycine, Lotus, and Medicago) and one non-legume (Arabidopsis). This will facilitate the integration of existing knowledge in nodule development with the *Arachis* transcriptome. The available data can be retrieved using a single or batch query or searching using gene ID, from above mentioned five species. The output displays the gene expression pattern in graphical as well as tabular form, along with further options to download the sequence data. The database is linked with PeanutBase, the main genomic resource of peanut. Additionally, the expression level of different splicing variants can be retrieved from the database. In summary, AhNGE serves as an important resource for the scientific community working on nodule development.

## Introduction

By virtue of their ability to fix (reduce) atmospheric di-nitrogen symbiotically, legumes are key to sustainable agriculture, providing a free and renewable source of reactive nitrogen to soils, crops and pasture systems. The legume-rhizobia symbiosis takes place inside specialized organs called nodules that develop on legume roots.

Peanut is one of the most economically-important legumes. Cultivated peanut, *Arachis hypogaea*, is an allotetraploid legume that originated ~9400 years ago due to the hybridization of two diploid progenitor species, *A. duranensis* (the A sub-genome donor) and *A. ipaensis* (the B sub-genome donor) (Bertioli et al. 2016; Bertioli et al. 2009). *A. hypogaea* contains an AABB-type genome (2n = 4X = 40) with genome size of ~2.7 Gb. Arachis diversified from the other sequenced legumes (such as soybean, common bean, *Medicago truncatula* and *Lotus japonicus*) ~58 million years ago (MYA) (Lavin et al. 2005; Lavin et al. 2001). Genetic studies in model legumes over last twenty years generated enormous knowledge with respect to molecular mechanisms underlying nodule development and metabolism (Roy et al. 2019). The genus *Arachis* belongs to the Dalbergioid clade of papilionoid legumes. Legumes belonging to this clade are more basal in their divergence within the papilionoid family with diverse atypical nodule developmental features. In *Arachis* species, rhizobia enter roots intercellulary via developmental fissures, a pathway called ‘crack entry,’ unlike intracellular entry via epidermal root hair cells in “more advanced” legumes. Also, *Arachis* species produce ‘aschynomenoid nodules’ harbouring infected cells without interspersed non-infected cells, as opposed to mosaic infected and uninfected cells typical of nodules of most (Medicago, Lotus, Soybean etc) other legume species. Interestingly, nodule development in Arachis shows a mixture of so-called ancient features, such as ‘crack entry’ and advanced features, such as ‘bacteroid terminal differentiation’(Boogerd and van Rossum 1997; Oono and Denison 2010). In a recent genome-wide phylogenomic study, it has been identified that a vital gene ‘RHIZOBIUM-DIRECTED POLAR GROWTH’ (*RPG*) required for nodule development is absent in the peanut genome, but present in all the other species examined. This study hypothesized that peanut nodule development is in an ‘intermediate step’ and might be on the verge of losing symbiosis (Griesmann et al. 2018). Though Arachis rhizobium infection/accomodation might be considered less advanced than that of other legumes in an evolutionary sense, yet the terminal bacteroid differentiation and highly effective SNF suggest advanced developmental features in Arachis (Bertioli et al. 2009; Janila et al. 2013). Research on this atypical but important mode of nodule development has taken little advantage of modern genomic tools (Das et al. 2019; Gupta et al. 2019; Karmakar et al. 2019; Kundu and DasGupta 2018; Sinharoy and DasGupta 2009; Sinharoy et al. 2009), mainly due to lack of a sequenced reference genome.

As a general feature, Dalbergia legumes, which are closely related to the Pterocarpus clade that includes *Arachis* species, exhibit ‘crack entry’ of rhizobia and develop terminally-differentiated bacteroids within nodules. In recent years, efforts have been made to establish *Aeschynomene evenia*, a legume belonging to the Dalbergia clade, as a model to study atypical nodule development (Chaintreuil et al. 2013; Czernic et al. 2015; Fabre et al. 2015; Lamouche et al. 2019a; Lamouche et al. 2019b). *A. hypogaea* belongs to the Pterocarpus clade, asister clade ofDalbergia. These two clades diverged ~49 MYA (Chaintreuil et al. 2013; Lavin et al. 2005). Transcriptome analysis has revealed differences in the expression of key genes during nodule development in *A. evenia* and *A. hypogaea*. These include Exopolysaccharide Receptor (EPR3), and a Flotillin-like protein (MtFLOT2/4) required for ‘root hair’ mediated infection in model legumes, which are highly-expressed in *A. hypogaea* but not in *A. evenia* (Karmakar et al., 2019; Gully et al., 2018). Hence, there is clear evolutionary divergence between Dalbergia lineages as far as nodule development ia concerned. Taken together, divergence of Arachis from *Aeschynomene sp*., recent polyploidization, the existence of the nodule developmental plasticity, and their economic value place peanut in an important position. A comprehensive understanding of peanut nodule development will be beneficial from evolutionary as well as economic points of view.

A database of spatiotemporal gene expression profiles of a model organism is an important community resource for novel gene function discovery and reverse genetics. With such a resource, a scientist does not need to perform RNAseq again and again. To integrate the existing knowledge from the last twenty years of genetic discovery in SNF and to apply it to Arachis, we have developed a database *‘Arachis hypogaea* Nodule Developmental Gene Expression’ (AhNGE). We conducted an RNA sequencing experiment on different time points during nodule developmental of *A. hypogaea* cv *Tifrunner* (the variety used for genome sequencing) (Bertioli et al. 2019b). The results were collated in a publicly-accessible, user-friendly, operating system (OS)-agnostic application, AhNGE. AhNGE enables users the following operations: a) individual gene-specific RNAseq data in graphical and tabular format; b) ortholog information of *Medicago truncatula* (both v4 and v5), *Lotus japonicus* (v3.0), *Glycine max* (v1) and *Cicer arietinum* (CDC Frontier and ICC 4958), and non-legume *Arabidopsis thaliana* (v11) obtained using Orthofinder program; c) novel transcript discovery table obtained from our experiments; d) batch downloading facility for *Arachis* CDS sequence (*A. hypogaea, A. duranensis* and *A. ipaensis*) and our transcriptome data; e) batch downloading facility for protein sequences of legumes whose ortholog information is available in the AhNGE database; and f) Basic Local Alignment Search tool (BLAST) interface to search the RNAseq data. AhNGE contains a search function where a particular gene (or batch), transcript ID, or a sequence can be input as a query. Moreover, the search function will accept gene IDs from other legumes (or even non-legumes), and display gene expression data of their Arachis orthologs. PeanutBase (http://peanutbase.org), the genomic resource of peanut is an excellent web interface for genomic research (Clevenger et al. 2016; Dash et al. 2016). From the tabular format of gene-specific RNAseq data available in the AhNGE database, one can directly connect to PeanutBase to obtain the genomic information available in the portal. Taken together, this database will be a powerful resourse for studying root-nodule development in peanut.

## Results

### Data generation

Five different time points were chosen for RNAseq transcriptome analysis to get a dynamic view of peanut nodule development. The time points were i), multicellular rosette-hair containing nodulation-susceptible root region (0 days post-inoculation, dpi) (Figure 1A), ii), primordium-forming susceptible zone (6 dpi) (Figure 1B), iii), nodules with dividing cells and differentiating bacteroids (10 dpi) (Figure 1C), iv), differentiated but weakly nitrogen-fixing nodules (15 dpi) (Figure 1D) and v), mature highly nitrogen-fixing nodules (21 dpi) (Figure 1E). An additional time point was also included to follow nodule senescence and capture senescence-related genes by applying full nitrogen (6 mM ammonium nitrate) at 19 dpi and harvesting nodules at 21 dpi, 48 hours after nitrogen treatment (21 dpi + NH_4_NO_3_). *A. hypogaea* cultivar Tifrunner seeds and *Bradyrhizobium sp. SEMIA 6144*, a strain that is widely used by the peanut community and secretes nod factor were used for the nodule developmental studies (Bertioli et al., 2019, Ibáñez and Fabra, 2011). RNAs are harvested from three independent replicates at six time-points. The detail of the RNAseq data and the statistics of the mapped reads in three independent Arachis genomes are given in Supplemental Figure 1.

**Figure 1:**
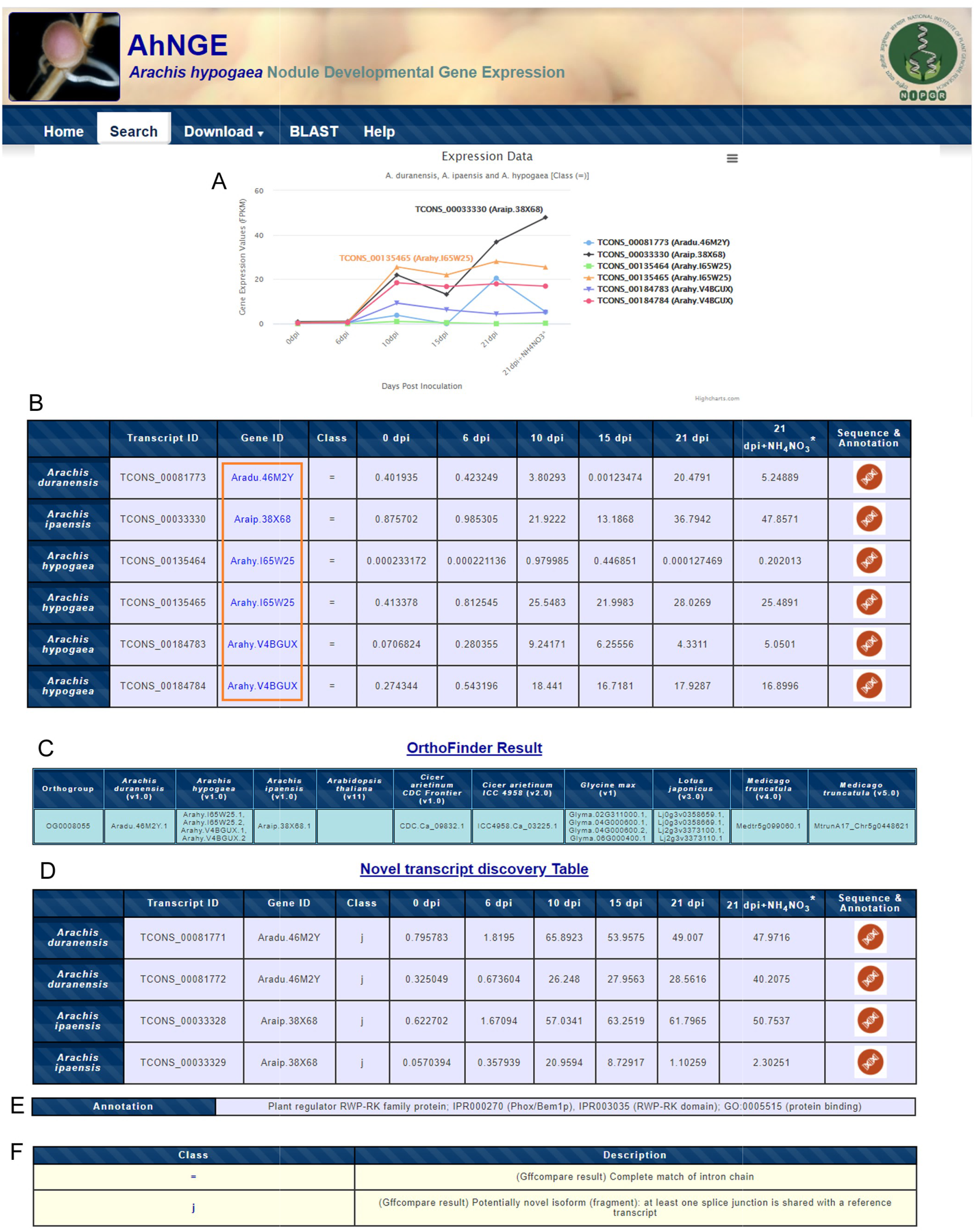
Macroscopic view of the progression of nodule development in Arachis. Root and rossets of root hairs at 0 dpi in the susceptible zone (A-B), (A) emergence of root hair from scar site (B) emergence of root hair from lateral root junction. Nodule primordia at 6 dpi (C-D), nodule primordia emergence on tap root (C), nodule primordia emergence from the lateral root junction (D), white non-nitrogen fixing nodule 10 dpi (E), pink nitrogen-fixing nodule (F) 15 dpi, and (G) 21 dpi. Scale bar = 0.5 mm.

### Orthology analysis

To understand the evolution of the nodule developmental program and to integrate the existing genetic knowledge and decipher the orthologous relationships among genes, OrthoFinder (Emms and Kelly 2015, 2019) was used. OrthoFinder is a robust platform for comparative genomics. By using rooted gene trees, it generates orthogroups and orthologs and identifies all the gene duplication events in those gene trees. This analysis is especially useful to identify gene duplication events. Proteomes of *A. duranensis* (v1.0), *A. ipaensis* (v1.0), *A. hypogaea* (v1.0), were retrieved from PeanutBase; *Arabidopsis thaliana* from TAIR (https://www.arabidopsis.org/); *Cicer arietinum CDC Frontier* (v1.0) (kabuli type chickpea), *Cicer arietinum* (ICC 4958 (v2.0) (desi type chickpea) from LIS-Legume Information System (https://legumeinfo.org/); *Glycine max* (v1) from Phytozome v12.1 (https://phytozome.jgi.doe.gov/pz/portal.html); *Lotus japonicus* (v3.0) from *Lotus* Base (https://lotus.au.dk/); *Medicago truncatula* (v4.0) from Medicago Genome Database (http://www.medicagogenome.org/); and *Medicago truncatula* (v5.0) from INRA Medicago Bioinformatics Resources (https://medicago.toulouse.inra.fr/MtrunA17r5.0-ANR/). The above-mentioned proteomes were used for orthogroup analysis using OrthoFinder-2.2.7 tool (Emms and Kelly 2015, 2019). The OrthoFinder analysis resulted in 36397 orthogroups, which are available in the database.

### Database design

The homepage of AhNGE (http://nipgr.ac.in/AhNGE/index.php) contains five major tabs. These are home, search, download, blast, and help. The ‘Home’ tab contains the project background, contact, and funding information.

#### Search tab

A user can feed a query through either the ‘Search’ tab or the ‘Blast’ tab. Multiple formats of gene/transcript ID are accepted in the search tab so that the users can initiate a search from a broad spectrum of legumes like chickpea, *Medicago*, or *Lotus*. This enables the user to understand the gene expression pattern of the peanut ortholog of a gene in question in other legumes. The search tab accepts gene IDs from any peanut sp. (*A. hypogaea* v1.0, *A. duranensis* v1.0 and *A. ipaensis* v1.0), *Medicago* (v4 or v5), chickpea (CDC Frontier v1.0 and ICC 4958 v2.0), soybean (v1), *Lotus* (v3) and *Arabidopsis* (v11).

If the query (or its peanut ortholog) is indeed expressed during nodule development in peanut, a multisection output page opens after search (Figure 2). The output page contains a graphical representation of the Transcripts (isoforms) expression pattern denoted with a ‘TCONS_’ prefix (Figure 2A), followed by a table of the RNAseq data in Fragments Per Kilobase Million (FPKM) values (Figure 2B). The first two sections only contain expression data for transcripts exactly matching with the primary transcript in the *Arachis* genome database PeanutBase (corresponding to the ‘=‘ form according to gifcompare result) (Figure 2A and B). These two sections display data from all the orthologs of the gene in question. Users can access the secondary data such as sequence and annotation by clicking the hyperlink present in the table (Figure 2B). In this table, the gene IDs are linked with hyperlink and by clicking those hyperlinks the user can directly go to PeanutBase feature page and will able to obtain the gene model and additional information on that page. The next table in the output page contains the OrthoFinder result which displays all the genes present in a particular orthogroup (Figure 2C). For instance, we have used the well-known transcription factor *Nodule Inception* (*NIN*) that controls nodule development (Marsh et al. 2007; Schauser et al. 1998), as a query. Orthologs/recent paralogs of *NIN* are present in all legumes but not in Arabidopsis. A user can retrieve the gene IDs and sequences of *NIN* from any legume (in the list) by feeding the known ID of Medicago or Lotus *NIN* as the query. This OrthoFinder table saves valuable time since the user does not need to run any phylogenetic analysis. Such an ortholog/recent paralog search function is available elsewhere (Legume Graph-Oriented Organizer—LeGOO) (Carrère et al. 2019), but neither Arachis nor chickpea is included in that analysis. The next section in the output page displays novel transcripts (Figure 2D). This table displays the gene expression values of the novel alternate spliced transcripts that were not annotated during the original release of the *Arachis* genome but were discovered during our RNAseq experiment. Under the novel isoform category, we only considered transcripts which at least have one splice junction shared with the reference genome. Many of the trancripts belong to this category shows over 10-fold induction in nodule indicating they are bonafied novel transcript. In the table 2D for *Araip.38X68* the novel transcript shows 102 fold upregulation at 15 dpi. This table helps the user to discover potentially novel functions of known genes. This is particularly useful since *Arachis* shows many atypical features of nodule development and morphogenesis (discussed in detail in the Introduction). The sequence of individual transcripts can be downloaded from both tables (Figure 2B and D) by clicking on the hyperlink. The annotation and the class code description of the transcript are available at the bottom of the table (Figure 2E and F). If the query gene does not express in any of the given conditions, the output page will only show the OrthoFinder table. Finally, if the query sequence is neither expressed during nodule development nor has an orthologue in other legumes, the output page will display ‘result not found’.

**Figure 2:**
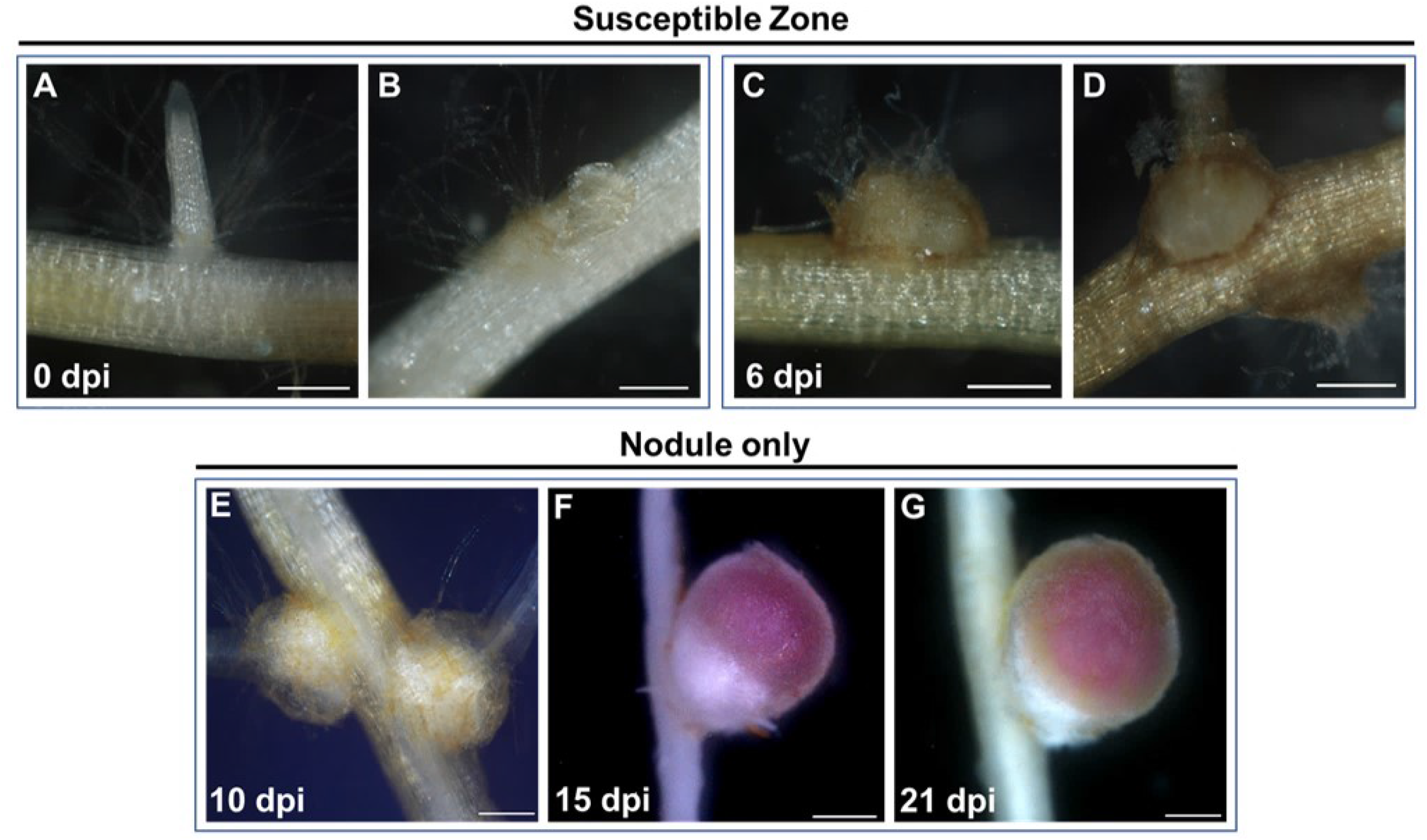
Screenshot of a result obtained after a search using Medtr5g099060.1 (*NIN*), an example: A, graphical representation of the transcript expression. B, tabular format containing gene expression data of the primary transcript. C, the ortholog information of six species (Arachis, Arabidopsis, Cicer, Glycine, Lotus, Medicago) D, annotation of the associated gene F, transcript class code information. The orange box highlighted the gene having hyperlinked and can be connected to PeanutBase.

#### BLAST tab

Users can also enter a query through the ‘BLAST’ tab. This search function accepts sequence information instead of gene or transcript IDs. Pairwise alignment is run against the assembled FASTA sequences. BLAST can be performed either against an individual *Arachis* genome, or against all three species.

#### Download tab

While the search tab is useful for a single entry of query, the ‘Download’ tab is useful for batch entry. Nonetheless, the download function can also be used to enter a single query sequence if the gene/transcript is from *Arachis*. This tab contains three options in the dropdown menu. In the first option, nucleotide sequence (FASTA) or expression values (Excel file) can be downloaded by providing gene ID(s). This section only accepts gene IDs and does not provide data for novel transcripts (see above). Data associated with novel transcripts discovered in this experiment can be retrieved by entering a query in the next section. The second section only accepts exact transcript IDs. In this section, careful interpretation of the output data is necessary. During the construction of the dataset, the reads were mapped to three different species of *Arachis* independently. But in each case, the transcript IDs generated were given continuous numbers like ‘TCONS_00000001 and TCONS_0000002’. Therefore, there is a chance that the same transcript ID may fetch data from different genes from the three species in question. The user can choose to download the expression value obtained by mapping against *A. duranensis, A. ipaensis, or A. hypogaea* from the option given at the bottom. The second section is more useful if a user wants to know the expression values of novel mRNA isoforms arising due to alternate splicing. The third download menu will allow a user to download protein sequences for all the species used in this database.

As stated above, AhNGE is a web server where the users can retrieve the gene expression data of any nodule-expressed peanut gene. Users can also search with a gene from other legumes, to determine whether the peanut ortholog is expressed during nodule development. The tetraploid cultivated peanut (*A. hypogaea*) genome sequence has been published following the sequencing of the parental genotypes (diploid) *A. duranensis and A ipaensis* (Bertioli et al. 2016; Bertioli et al. 2019b). Keeping the complexity of the allotetraploid genome in mind, we mapped the RNAseq data of *A. hypogaea cv* Tifrunner to both progenitor diploid genomes, and as well as the newly sequenced allotetraploid AABB genome (Bertioli et al. 2016; Bertioli et al. 2019a).

## Discussion

AhNGE is created to streamline comparative functional genomic and transcriptomic studies on peanut nodule development. How did the root nodule symbiosis evolve in different phylogenetic clades from a single hypothetical point of predisposition event remains an open question (Griesmann et al. 2018; Raul et al. 2019). To obtain the answer, thorough investigation is needed across diverse phylogenetic taxa focusing on nodule development. After two decades of genetic research, more than 200 genes have been identified as vital players in nodule development (Roy et al. 2019). The function, regulation, and genetic variation present in orthologs of these genes in crop legumes will help to answer not only evolutionary questions, but also will be an avenue towards breeding to improve SNF in legumes. With next-generation sequencing now common, obtaining large-scale data for functional genomics is no longer a bottleneck, although analysis of such data can be. This website shunts the need of users to download large datasets of individual legumes from web servers like Legume information system (LIS, https://legumeinfo.org/species), followed by a synteny analysis or phylogeny analysis. AhNGE has all the orthogroups, already analyzed. This covers mostof the relevant legumes and one non-legume (Arabidopsis).

Peanuts belong to the Dalbergioid clade with many atypical nodule developmental features (Boogerd and van Rossum 1997; Sinharoy and DasGupta 2009; Sinharoy et al. 2009). It’s divergence from model legumes and from its closely studied relative, *Aeschynomene sp*, happened around 50 MYA (Chaintreuil et al. 2013; Lavin et al. 2001). Feas liketure the formation of multicellular root hair cells and crack invasion of symbionts, absence of the ortholog of *RPG* from the peanut genome, and symbiont promiscuity suggest that this plant is in a primitive state in the evolution of nodule infection/entry, and might be on the verge of losing symbiosis (Boogerd and van Rossum 1997; Griesmann et al. 2018; Tajima et al. 2008). Whereas, terminal bacteroid differentiation, absence of *NCR* genes from peanut nodule transcriptome, high nitrogen fixation efficiency, and worldwide peanut cultivation without any nitrogen fertilizer application independent evolution of this clade in the last 50 MYs which helped it to acquire more modern quality traits of nodule development (Janila et al. 2013; Oono and Denison 2010; Raul et al. 2019). Recent sequencing of peanut genomes has created opportunities to study genetic details of nodulation in this family. This database contains six RNAseq data points from different stages of peanut nodule development analyzed against all three peanut genomes, with user-friendly data retrieval system containing 36397 orthogroups’ expression. To conclude, as an important community resource, AhNGE has the potential to serve as a platform for dissecting the intriguing and atypical features of nodule development in peanut.

Although AhNGE at present has six RNAseq datasets, it will be updated from time to time with more RNAseq data from peanut nodule knockdown/knockout mutants, ecotypes, and different nodule developmental stages. Moreover, inclusion of a gene regulatory network after the integration of these additional datasets will further accelerate comparative genomic studies through this portal.

## Material and Methods

### Sample collection and RNA sequencing

Peanut seed pods were treated 5 min with 30% Clorox and further rinse with water. The seeds were then coated with Vitavex for 4 Hours and platted onto a wet filter paper in dark at 26-28°C. After four days the seeds started germinating at this point, they were saw on thoroughly washed, twice autoclaved 2:1 turface: vermiculite in 28-30°C in a 10-inch pot. Plants were cultivated under a 14-h/10-h light/dark regime with 200 μE. m^-2^. s^-1^ light irradiance at 28/22°C (day/night) and 40% relative humidity and were inoculated with Bradyrhizobium SEMIA6144 (grown in TY broth at 28°C), 12 days post potting. For inoculation bacterial pellets were suspended in B&D without nitrogen (R) to an OD of 0.02 - 0.04 and 50 ml were added to each pot after which plants were bottom watered. This experimental were repeated three times with freashly grown bacteria but under identical growth condition. Time points for harvest of tissues were 0, 6, 10, 15, 21 dpi and 19 dpi + 2 days in presence of 6 mM nitrogen (designated as 21 + NH_4_NO_3_). For 0 and 6 dpi, the susceptible zone was harvested (root tips were eliminated), and from 10 dpi onward nodules were harvested. Total RNA was extracted from three independent biological replicates using TRIZOL reagent (Life Technologies), followed by genomic DNA removal by DNase 1 (Ambion), and additional column purification with RNeasy MinElute CleanUp kit (Qiagen). The RNA obtained in this way have RIN number between 8.9 – 10 obtained using Agilent 2100 Bioanalyzer. Kapa stranded RNA-seq kits (Kapa Bio Systems) was used to generate paired-end libraries at the Georgia Genomics Facility (Athens, Georgia).

### Transcript assembly and quantification

De-multiplexed paired-end reads were checked for quality using FastQC (http://www.bioinformatics.babraham.ac.uk/projects/fastqc/) including base composition and kmer overrepresentation. After inspection, 10 bases from the 5’ end of each read were trimmed due to nonrandom base composition. All reads maintained high quality (>30) over the total length of the read. After QC, reads were mapped to an rRNA database using Bowtie v 1.1.0 to remove any rRNA contamination. The RNAseq data then were analysed using the Cuffdiff v2.0.2 (Trapnell et al. 2012). The Cuffdiff suite analysis was carried out independently three times using *A. duranensis, A. ipaensis*, and *A. hypogaea* genomes. The following workflow were used for the analysis. Using Tophat v1.3.2 (Trapnell et al. 2012), the reads were aligned to the *A. duranensis, A. ipaensis*, and *A. hypogaea* cv Tifrunner genomes separately. The genomes and indexed reference files were downloaded from the Peanutbase. Tophat first used Bowtie v (R) for an initial alignment of the reads to the genomes and the rest of the reads were then aligned to exons to generate potential splice junctions. Cufflinks v2.0.2 (R) was used to process the Tophat alignments. By combining the sequencing data with the available genome information, Cufflink generated assemble transcript. Cufflink treated the regions of the genome that were covered by RNAsequence reads as potential exons. The abundance of assembled transcripts was calculated using fragments per kilobase of exon per million fragments mapped (FPKM) unit. The FPKM calculation taken care the normalization of the library’s sequencing depth during transcript abundance calculation across the nodule development time series. The annotation available in the Peanutbase was used for the reference annotation-based transcript assembly. During this analysis novel discoveries including new splice sites, transcriptional start sites were identified using Cuffcompare. The assembled Cufflinks transcripts were separately processed with both the Cuffmerge and Cuffcompare analytical strategies. The minimum set of transcripts identified by Cufflinks as best describing the aligned reads in each sample was first processed with Cuffcompare to classify the transcripts based upon existing annotation. The gffread utility tool was used to generate a FASTA file. The transcripts (isoforms) expression pattern was denoted with a ‘TCONS_’ prefix. The cleaned reads per library and the % of total mapped reads are shown in supplemental table 1.

### AhNGE web interface design

The AhNGE database was designed using the web interface queries and the Apache Hypertext Transfer Protocol (HTTP) server (Version: 2.2) to access the data from MySQL (Version: 8.0) database. MySQL is a relational database management system (RDBMS) based on Structured Query Language (SQL) that is used to store and manipulate the data in the database. Data retrieval from stored data was accomplished using MySQL commands. Hypertext Markup Language (HTML), Hypertext Pre-processor (PHP 7.2), Cascading Style Sheets (CSS) and Javascript are used on the front-end to develop the database and dynamic page. All development was carried out on a Linux OS (Ubuntu 16.04.4) platform. The graphical representation of the expression values at the web interface is visualized using Highchart cloud.

## Acknowledgments

The authors are thankful to DBT (Department of Biotechnology)-eLibrary Consortium (DeLCON), India for providing access to e-resources.

